# Microsatellite Density Landscapes Illustrate Short Tandem Repeats Aggregation in The Complete Reference Human Genome

**DOI:** 10.1101/2022.04.16.487617

**Authors:** Yun Xia, Douyue Li, Tingyi Chen, Saichao Pan, Hanrou Huang, Wenxiang Zhang, Yulin Liang, Yongzhuo Fu, Zhuli Peng, Hongxi Zhang, Liang Zhang, Shan Peng, Ruixue Shi, Xingxin He, Siqian Zhou, Weili Jiao, Xiangyan Zhao, Xiaolong Wu, Lan Zhou, Jingyu Zhou, Qingjian Ouyang, You Tian, Xiaoping Jiang, Yi Zhou, Shiying Tang, Junxiong Shen, Kazusato Ohshima, Zhongyang Tan

## Abstract

**Background:** Microsatellites are increasingly realized to have biological significance in human genome and health in past decades, the assembled complete reference sequence of human genome T2T-CHM13 brought great help for a comprehensive study of short tandem repeats in the human genome.

**Results:** Microsatellites density landscapes of all 24 chromosomes were built here for the first complete reference sequence of human genome T2T-CHM13. These landscapes showed that short tandem repeats (STRs) are prone to aggregate characteristically to form a large number of STRs density peaks. We classified 8,823 High Microsatellites Density Peaks (HMDPs), 35,257 Middle Microsatellites Density Peaks (MMDPs) and 199, 649 Low Microsatellites Density Peaks (LMDPs) on the 24 chromosomes; and also classified the motif types of every microsatellites density peak. These STRs density aggregation peaks are mainly composing of a single motif, and AT is the most dominant motif, followed by AATGG and CCATT motifs. And 514 genomic regions were characterized by microsatellite density feature in the full T2T-CHM13 genome.

**Conclusions:** These landscape maps exhibited that microsatellites aggregate in many genomic positions to form a large number of microsatellite density peaks with composing of mainly single motif type in the complete reference genome, indicating that the local microsatellites density varies enormously along the every chromosome of T2T-CHM13.

## 1. Background

Tandem repeat biology is causing a revolution in genetics in the past two decades as many studies of understanding the evolution and biological functions of tandem repeats in the genomes of thousand species and also human diseases [1–4]. Indeed, more than half of the human genome is constituted of repetitive sequences making them the most challenging genomic regions to study, and repetitive sequences are generally classified into interspersed repeats and tandem repeats [5]. The Short Tandem Repeats (STRs), also called microsatellites, happen ubiquitously in human genome scattered in coding and non-coding regions, which commonly occur with repeat units of 1-6 base pairs and own the highest mutational rate in genome [1, 3, 6]. Microsatellites are increasingly realized to have biological significance in human genome and health, and are reported to involve in more than 30 disorders and several cancers [7–10], regulate gene expression in healthy genomes [11, 12], and also be related to genetic plasticity and missing heritability [13–20]. Slipped-strand mispairing was suggested as a major mechanism for tandem repeat occurrence [21, 22], and we formerly presented a folded slippage model for short tandem repeats occurring mechanism, predicting that micro-disturbing in the process of replication may provide a lot of chances for the template chain folded to produce short tandem repeats, which are possibly selected and fixated for different biological significance in the long-history evolution [23].

Although short tandem repeats have been studied in human genome for several decades, many sequenced columns of human genome so far contain a large number of unsequenced gaps, and these unsequenced gaps are often composed of short tandem repeats [3, 5]. To date, studies about short tandem repeats on complete, gap-free human genomes have been very lacking. The recent assembly of T2T-CHM13 reference removes the gap filled regions of autosomes and Chromosome X [5, 24–28], and over 50% gaps in Chromosome Y [29], therefore it represents a truly complete human genome, this complete reference sequence of human genome T2T-CHM13 will certainly bring great helps for comprehensive study short tandem repeats in human genome. And so far, there is still no unified threshold for microsatellites studies, the repeats in animal genomes are generally considered to be longer than 12 bp [30, 31], and 3% of human genomes was reported consist of microsatellites under that standard [32]. And microsatellites of human sequences comprised of GA/TC/GC/AT bases are investigated using seq-requester microsatellite [24]. However, we applied a threshold of 6, 3, 3, 3, 3, 3 iterations for mono- to hexa-nucleotide repeat motifs for analyzing microsatellites in the complete reference human genome T2T-CHM13, the threshold was widely used to study microsatellites in the genome sequences of viruses, mitochondrial and chloroplast [2, 33–35]. And we have first applied the threshold to investigate microsatellites in the Y-DNA of the human reference genome (GRCh38, NC_000024.10) at 1 Kbp resolution by Differential Calculator of Microsatellite version 2.0 (DCM 2.0) method, revealing an exact distributional feature of STRs in every local bins of 1 Kbp sequence of the chromosome Y [36]. Herein, we built 24 Microsatellite landscape maps in all 24 chromosomes of the first complete human genome T2T-CHM13 (CHM13), including the positions and motif types of all classified High Microsatellite Density Peaks (HMDPs), Middle Microsatellite Density Peaks (MMDPs), and Low Microsatellite Density Peaks (LMDPs).

## 2. Materials and methods

### 2.1. Genome sequences

The complete 24 chromosome sequences of the human reference genome T2T-CHM13v2.0 were collected from GenBank, and the accession No. of the complete 24 chromosome sequences were listed in Table S2. All chromosome sequences of human reference genome GRCh38.p14 were also obtained from GenBank (Table S2).

### 2.2. STRs identification

The Imperfect Microsatellite Extractor (IMEx 2.1) [37] was applied to identify STRs in all the sequences of T2T-CHM13v2.0 and GRCh38.p14 genomes. The threshold for extracting perfect tandem repeats was set at iterations of 6, 3, 3, 3, 3, 3 for mono-, di-, tri-, tetra-, penta- and hexa-repeat motif respectively.

### 2.3. Calculate local microsatellite density at 1 kb resolution

The local microsatellite density of 24 complete chromosome sequences of T2T-CHM13 were counted by the new promoting program of Differential Calculator of Microsatellites version 3.0 (https://github.com/zhongyangtan/DCM.git), which calculate the local STRs density by divided the every chromosome sequence into a large number of differential-units (bins) with size of 1kb, so microsatellite position-related Differential-unit_1kb_ (D_1_) Relative Density (pD_1_RD) was calculated for every 1 kb sequence of the 24 chromosome sequences of T2T-CHM13, and also for GRCh38.p14, which was proven to be a better resolution to calculate local STRs density of human genomic sequence [36]. The calculation formula for pD_1_RD is:

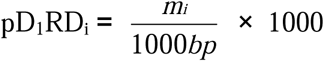

In this formula, pD_1_RD_i_ is the pD_1_RD in the i-th 1 kb bins of the genome, m_i_ is the size (bp) of microsatellite in the i-th 1 kb bins.

### 2.4. Identifying Microsatellite Density Peaks

After the microsatellite density in every 1 kb bins, i.e, pD_1_RD, of the genome was calculated. The adjacent bins with similar density ranges were merged, and each merged density bin is referred to as a peak. When the pD_1_RD values of each bin in adjacent bins is greater than or equal to 90 and less than 150 (150 < pD_1_RD ≥ 90), those adjacent bins was identify as a Low Microsatellite Density Peak (LMDP); the pD_1_RD values of each bin in adjacent bins is greater than or equal to 150 and less than 300 (300 > pD_1_RD ≥ 150), those adjacent bins was identified as a Middle Microsatellite Density Peak (MMDP); and the pD_1_RD values of each bin in adjacent bins is greater than or equal to 300 (pD_1_RD ≥ 300), those adjacent bins was identify as a High Microsatellite Density Peak (HMDP).

### 2.5. Building the STRs density landscape maps

A series of pD_1_RD values were obtained by T2T-CHM13 chromosome sequences by the DCM v3.0. Maps displayed the series of pD_1_RD values from telomere of p-arm to telomere of q-arm of every chromosome by using the UCSC Genome Browser’s bedGraph format. Then, 24 landscape maps of STRs density for T2T-CHM13 were built in the viewer of UCSC Genome Browser window, in which a full chromosomal view of local STRs density is shown for every bin of 1 kb sequence of the chromosome from telomere of p-arm to telomere of q-arm. 24 landscape maps of STRs density for GRCh38.p14 were also built.

### 2.6. Sorting Microsatellite Density Peaks

High Microsatellite Density Peaks (HMDPs) (pD_1_RD ≥ 300), Middle Microsatellite Density Peaks (MMDPs) (300 > pD_1_RD ≥ 150) and Low Microsatellite Density Peaks (LMDPs) (150 > pD_1_RD ≥ 90) were sorted by the new developed program Microsatellite Density Peaks Sorter version 1.0 (MDPS.v1.0, https://github.com/zhongyangtan/MDPS.git). And the MDPS.v1.0 also further sorted HMDPs, MMDPs and LMDPs into different Main Motif Type (MMT) and sub-Main Motif Type (sub-MMT). (Table S5, S6).

### 2.7. Mapping Microsatellite Density Peaks

The sorted data of exact position, corresponding motif type and peak name of HMDPs, MMDPs and LMDPs in every chromosome were transferred by the UCSC Genome Browser’s bed format and displayed under the landscapes with three tracks of HMDPs, MMDPs and LMDPs in the viewer of UCSC Genome Browser window.

### 2.8. Comparison of HMDPs between T2T-CHM13.v2.0 and GRCh38.p14

The similar HMDPs between T2T-CHM13.v2.0 and GRCh38.p14, were firstly determined allelic positionby MUMmer v3.23 [38], then comparing motif type by new developed program Genome HMDPs Comparator version 1.0 (GHC.v1.0, https://github.com/zhongyangtan/GHC.git), finally, aligned by ClustalX v2.1 [39].

### 2.9. The division of Genomic Regions

The full genome of T2T-CHM13 was divided into different Genome Regions according to local microsatellites density features (Table S9). The Genomic Regions with the extremum pD_1_RD value ≥ 300 between bins were classified as High Variable Microsatellite Density Region (HVMD-R), those with (300 < the extremum pD_1_RD value ≥ 150) between bins were classified as Middle Variable Microsatellite Density Region (MVMD-R), and those with (150 > the extremum pD_1_RD value ≥ 90) between bins were classified as Low Variable Microsatellite Density Region (LVMD-R) subclass. The Genomic Regions with the extremum pD_1_RD value < 90 between bins were classified as relatively Even microsatellite density (E-) Region class. Genomic Regions clustering with single dominant motif type of peaks were classified as Peak Cluster (PC-) Region class and Telomere Repeat (T-) Region class (located in the telomere regions).

## 3. Result

### 3.1. Microsatellite Density landscapes

The recently released complete assembles of T2T-CHM13 including all 22 autosomes, chromosome X and Y, comprise of 3,117,275,501 bp of DNA; and we obtained 22,198,470 microsatellites with total size of 183,133,984 bp and microsatellites relative density (RD) is 58.75 in the full genome (Table 1-a, Table S2 & S3). The widely used method of relative density for analyzing microsatellites was proved limited for analyzing STRs in big sequence like human genome; therefore, to explore the exact distributing feature, the microsatellites landscapes at 1 Kbp resolution, were formerly comprehensively surveyed in the Y-DNA of reference human GRCh38 by the differential calculator of microsatellites [36]. Herein, STRs of the complete reference human genome T2T-CHM13 including all chromosomes were investigated by the differential calculate method, in which the values of position related relative density at 1 kilo-base resolution (pD_1_RD) were calculated in every bin unit of 1kb DNA sequence, and the pD_1_RD value was suggested to be possibly a better way to estimate the local microsatellite density variation in human genome [36]. The bedgraph format tool of UCSC Genome Browser was used to map the exact STRs distribution features with the pD_1_RD data of the full T2T-CHM13 genome, a series of panoramic landscapes of microsatellites density were obtained to precisely exhibit the local microsatellites density in every bin of 1 kb sequence for the 22 autosome, chromosome X and Y of T2T-CHM13 in viewers of Genome browser window (Fig. 1, Fig. 2, Fig. S1, Fig. S2, Table S1). The 24 landscapes revealed that the relative density in every 1 kb sequence (the pD_1_RD value) varies enormously in different site along the 24 chromosomes even from 0 to 970, and the STRs is prone to accumulate to a large numbers of different extent microsatellite density peaks; these density peaks are found genome-wide, and were classified into 8,823 high microsatellites density peaks (HMDPs) (pD_1_RD≧300), 35,257 middle microsatellites density peaks (MMDPs) (300>pD_1_RD≧150) and 199, 649 low microsatellites density peaks (LMDPs) (150>pD_1_RD≧90) (Table 1-b, Table S4.01-S4.06); and the bed format tool of UCSC Genome Browser was used to display the exact position and motif types of these HMDPs, MMDPs and LMDPs (Fig. 2, Fig. S2). Similarly, the microsatellites landscapes were comprehensively surveyed in the reference human genome GRCh38 as comparison (Table 1-b, Table S14 & S15, Fig. S12).

**Figure 1.**
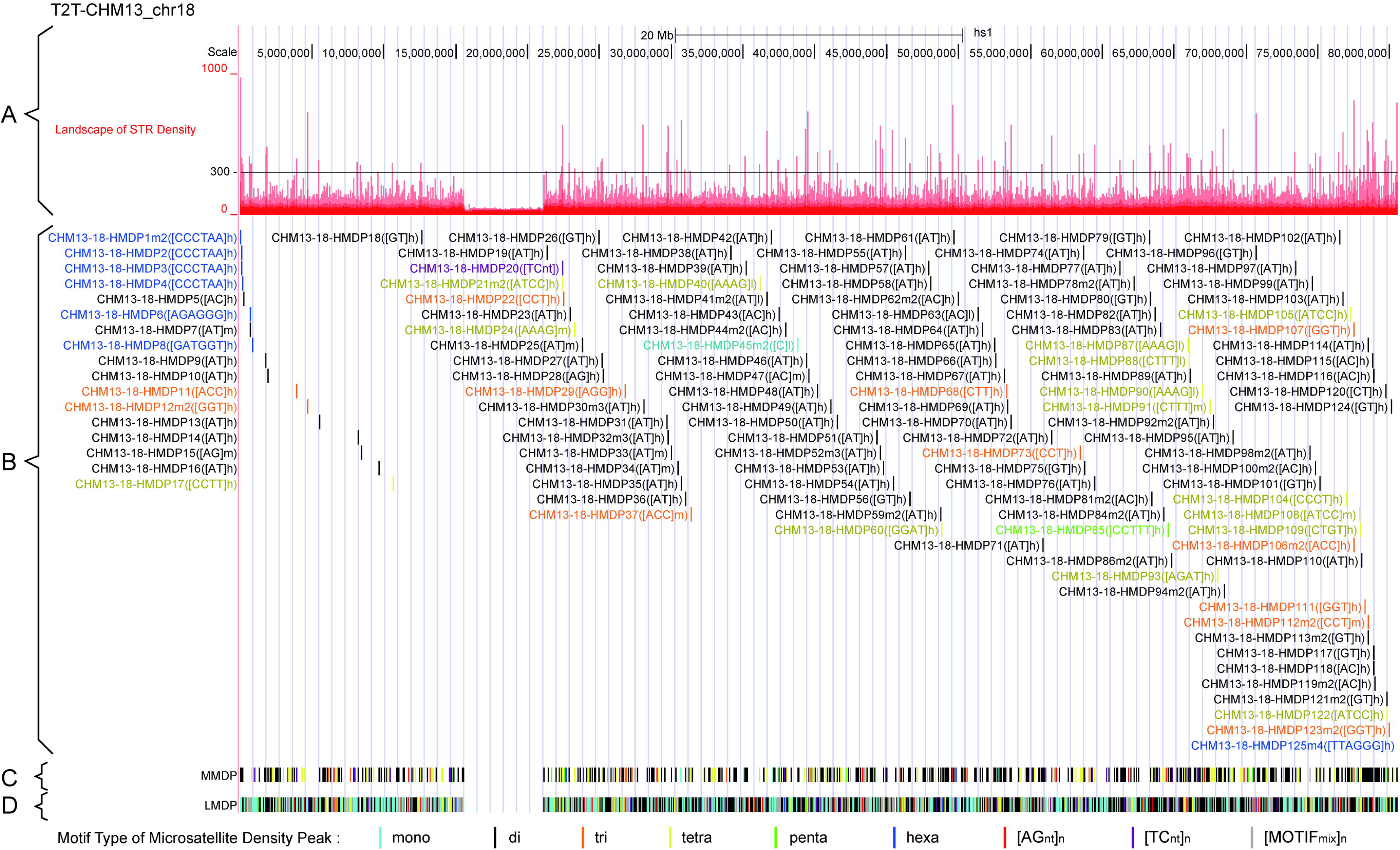
The landscape of STR Density, HMDP (High Microsatellite Density Peak), MMDP (Middle Microsatellite Density Peak) and LMDP (Low Microsatellite Density Peak) in T2T-CHM13 chromosome 18 with smallest HMDPs. **A.** Landscape of STR Density is displayed on the first track, and the microsatellite density values are shown on the ordinate. **B.** HMDPs are displayed in pack mode on the second track, including its Motif Type, location (shown in colored vertical line) and name. The HMDP name consists of “Genome name”, “chr No.”, “HMDP No.”, “HMDP size” and “sub-Main Motif Type (sub-MMT)”. **C-D.** MMDPs and LMDPs are displayed in dense mode on the third and last track. Chromosome 18 with the lowest number of HMDP, is shown here, and other chromosomes of the T2T-CHM13 are shown in Figure S2.

**Figure 2.**
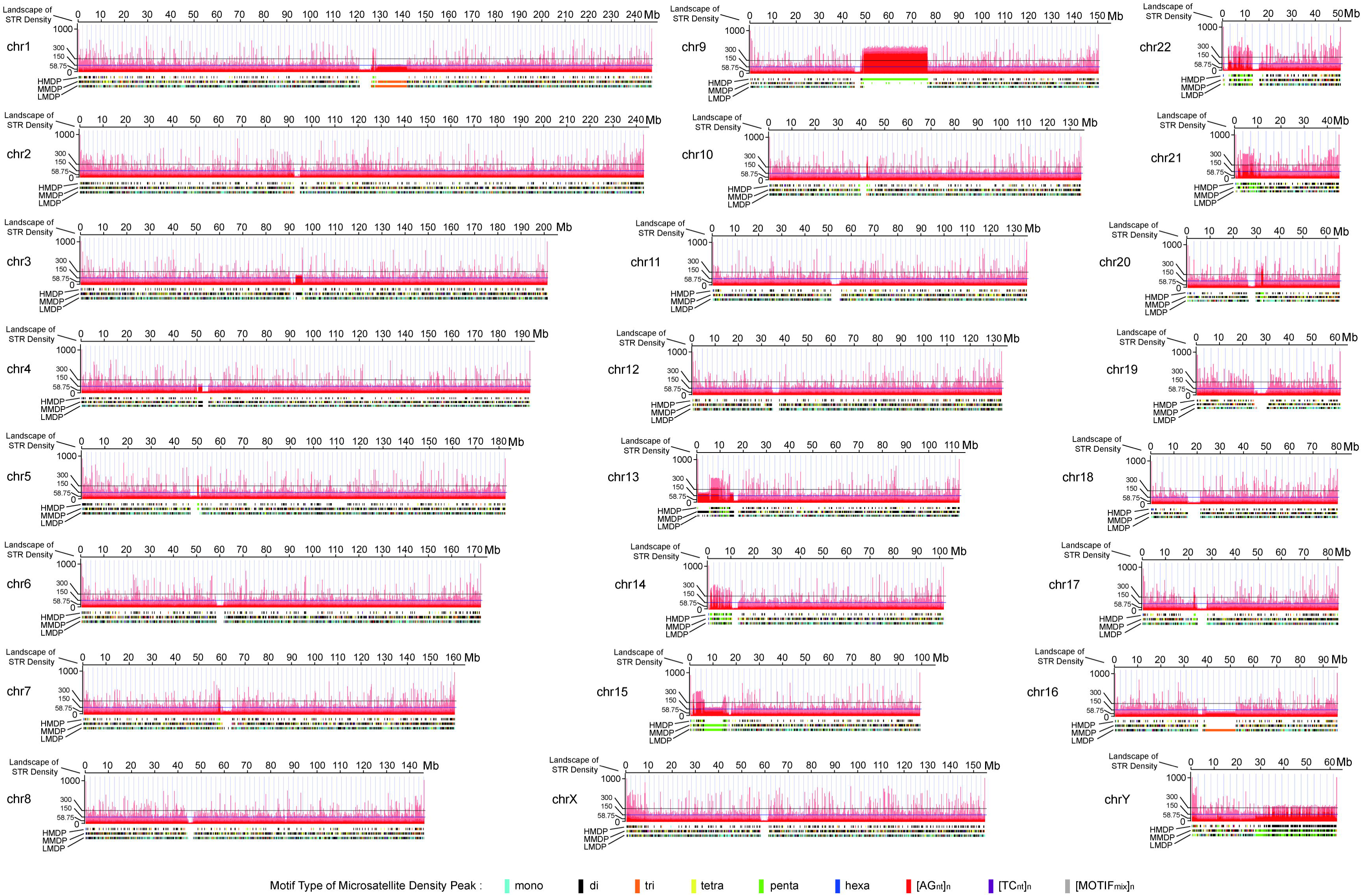
Landscape map of STR Density of 24 chromosomes of T2T-CHM13. Landscape of each chromosome, first track displayed landscape of STR Density, second track displayed HMDP, third track displayed MMDP, last track displayed LMDP.

**Table 1.** Statistic of microsatellite density landscape information on the human genome T2T-CHM13. **a.** The assembly information and microsatellite statistics of T2T-CHM13v2.0 and GRCh38.p14. **b.** The list of microsatellites density peaks in T2T-CHM13v2.0 and GRCh38.p14. **c.** The motif types of HMDP in T2T-CHM13v2.0. **d.** Genomic regions characterized by microsatellite density in T2T-CHM13v2.0.

### 3.2. High Microsatellite Density Peaks (HMDPs)

The local relative density values (pD_1_RD≧300) of those HMDPs are approximately six times or more of the average relative density (RD=58.75) of the full T2T-CHM13 genome, the significant statistical bias of microsatellites relative density in local genome region hints their importance to human genome (Fig. 3-A, Fig. S2.01). It was found that all these HMDPs occur genome widely in all the 24 chromosomes of the T2T-CHM13, especially pervade in the two arms of all chromosomes at different intervals, except absent at most centromeric region and some pericentromeiric region. The largest quantity of HMDPs was found in ChrY with 2,697 HMDPs, secondly in Chr13, the minimum number of HMDPs in Chr19 with 127 HMDPs identified, and only 305 HMDPs were found on chr1 with longest sequence; therefore, the distribution of HMDPs is not related to chromosome size directly. Furthermore, 89.26% of the HMDPs are identified in intergenic, 10.74% in intron region, but no HMDPs in exon region (Fig. 3-B-a, Table S10.01).

**Figure 3.**
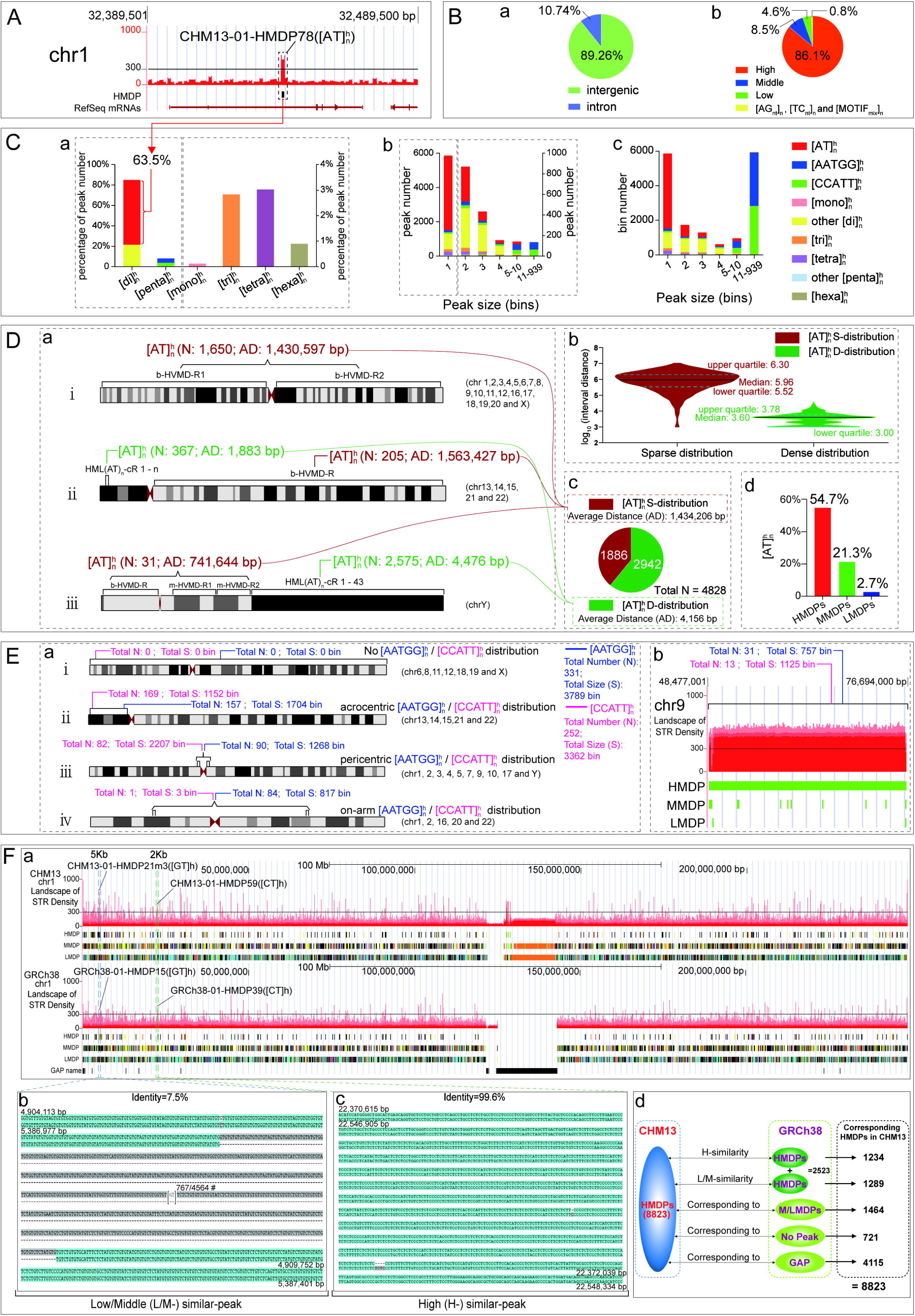
Summary of HMDPs in T2T-CHM13. **A.** An [AT]_n_^h^ HMDP in the T2T-CHM13. **B.** (a) Ratio of HMDP corresponding to intron and intergenic regions. (b) Ratio of hMMT, mMMT and lMMT HMDPs and other HMDPs. **C.** Statistic of HMDP number and bin, (a) The percentage of peak number of mono-, di-, tri-, tetra-, penta- and hexa-hMMT HMDPs in all hMMT HMDPs; (b, c) The number and total bin number of hMMT HMDP in each Peak sequence size (bins) group. **D.** [AT]_n_^h^ HMDP distribution, (a) The 3 [AT]_n_^h^ HMDP distribution models; (b) The distribution map of interval distance of [AT]_n_^h^ Sparse distribution (S-distribution) and Dense distribution (D-distribution) HMDPs, the ordinate indicated the logarithm base 10 of the interval distance (bp), and the values of the interval distance are listed in Table S10.03; (c) The number of [AT]_n_^h^ Sparse distribution and Dense distribution HMDPs; (d) The percentage of [AT]_n_^h^ HMDPs, MMDPs and LMDPs to all HMDPs, MMDPs and LMDPs. **E.** [AATGG]_n_**^h^** and [CCATT]_n_**^h^** HMDP distribution, (a) The 4 distributional models of [CCATT]_n_^h^ and [AATGG]_n_^h^ HMDPs; (b) A big cluster of [AATGG]_n_**^h^** and [CCATT]_n_^h^ motif type HMDPs at the p-arm of Chr9. **F.** Comparison of landscapes T2T-CHM13.v2.0 and GRCh38.p14, (a) Comparison in Chr 1; (b) identity of a pair low similar HMDPs; (c) identity of a pair high similar HMDPs; (d) corresponding HMDPs of the T2T-CHM13 corresponding to MDPs of GRCh38.p14.

### 3.3. Main Motif Types (MMTs) of HMDPs

Analysis of the composition of the motifs in every HMDP demonstrated that same type of motif is prone to accumulate in the same HMDP, and therefore, we classified these HMDPs by the main motif composition in each HMDP; for example, when the AT motif account for the main motif composition in a HMDP, it was named (AT)_n_ Main Motif Type (MMT) HMDP (Table S4.). And the MMT HMDPs can also be further classified into three sub-types: the high-percentage Main Motif Type (hMMT) HMDPs (main motif ≧ 66.7%), the middle-percentage Main Motif Type (mMMT) HMDPs (66.7% > main motif ≧ 50%), and the low-percentage Main Motif Type (lMMT) HMDPs (50% > main motif ≧ 33.4%) (Table 1-c, Table S4-S8).

All 8,823 HMDPs be classified into Main Motif Type (MMT) class and Mix Motif Type class. Though the microsatellites generally comprising tandem repeats with motifs of 1-6 bp makes that there are possible total 964 motif types (Table S16), we actually identified only 87 MMTs from 8819 HMDPs belong to MMT class, and these 87 MMTs were further classified into 8 subclasses: mono-, di-, tri-, tetra-, penta-, hexa-, AG- and TC-MMT subclass. The di-MMT subclass includes 6944 HMDPs is the most abundant HMDPs subclass, secondly is the penta-MMT subclass. Notably, 86.1% of the total 8823 HMDPs are the hMMT HMDPs (Fig. 3-B-b,Table S10.01), suggesting that high percentage of same repeat motif is prone to accumulated into same high microsatellites density peak (HMDP) in the complete reference human genome T2T-CHM13. Moreover, many MMT HMDPs usually appears in pairs with motif reverse complementary in the complete genome like the example of 386 (AATGG)_n_ MMT HMDPs reversely complementary pairing with 311 (CCATT)_n_ MMT HMDPs (Table 1-c).

### 3.4. [AT]_n_^h^ motifs type are the most abundant HMDPs

Among the all hMMT HMDPs, 4828 HMDPs are [AT]_n_^h^ motif type, representing 63.5% of the di-MMT subclass HMDPs and also 54.7% of all 8823 HMDPs, therefore, it is the most abundant hMMT HMDPs (Table 1-c, Fig. 3-C, Table S10.02). These [AT]_n_^h^ HMDPs were classified into two distributional patterns by average distance (AD): sparse-distributional (AD: 1,434,206 bp) and dense-distributional (AD: 4,476 bp) patterns, there are 1886 [AT]_n_^h^ HMDPs scatter sparsely and 2942 [AT]_n_^h^ HMDPs array densely in the full T2T-CHM13 genome (Fig. 3-D-a,b,c, Table S10.03, Fig. S3.01). As a whole, the distributional features can be summed up to 3 [AT]_n_^h^ HMDPs chromosomal distribution models, one model is that only [AT]_n_^h^ HMDPs sparse-distribution appear in the two arms of the 19 chromosomes, another is that [AT]_n_^h^ HMDPs dense-distribution occur in the short p-arm and sparse-distribution in the long q-arm as found in the five acrocentric chromosomes, the other is the [AT]_n_^h^ HMDPs distribution in the Chr Y which 35 [AT]_n_^h^ HMDPs with sparse-distribution are found in Chr Y euchromatin region and 2575 [AT]_n_^h^ HMDPs with dense-distributions in Chr Y heterochromatin region. However, the [AT]_n_^h^ HMDPs is abundantly representing 54.7% of all HMDPs, but 21.3% of all MMDPs and only 2.7% of all LMDPs (Fig. 3-D-d), suggesting that AT motif tandem repeats is easy to accumulate at high density in the complete reference genome. And (AT)_n_ repeats aggregation are reported to related to chromosomal structure and rearrangements [40–42].

### 3.5. The [AATGG]_n_^h^ and [CCATT]_n_^h^ penta-hMMT HMDPs account for about half sequence size of all hMMT HMDPs

There are 773 penta-motif HMDPs identified in the full T2T-CHM13 genome, and ranks secondly in all 8 subclasses of the MMT HMDP class; however, the reverse complementary [AATGG]_n_^h^ and [CCATT]_n_^h^ hMMT HMDPs are found 331 and 252 in numbers accounting for 75.4% of all penta-hMMT HMDPs (Table 1-C, Table S4 & S5). Though the large majority of the two types are small in size that is lower than 5 bins (5 kbp); the HMDPs with size that is larger than 5 bin, are mainly [AATGG]_n_^h^ and [CCATT]_n_^h^ hMMT HMDPs (Fig. 3-C-b & c, Table S10.02); therefore, total size of [AATGG]_n_^h^ and [CCATT]_n_^h^ hMMT HMDPs are 3789 and 3362 bins respectively, and combination of two hMMT HMDPs accounting for about 43.55% of the size of all hMMT HMDPs, implied their importance to human genome structure. Analysis of occurred locations of the [AATGG]_n_^h^ and [CCATT]_n_^h^ hMMT HMDPs reveals 4 distributional models about the two hMMT HMDPs in the T2T-CHM13 genome: (a) no [AATGG]_n_^h^ and [CCATT]_n_^h^ hMMT HMDPs distribution occur in Chr 6, 8, 11, 12, 18, 19 and X, (b) both the two hMMT HMDPs appear in short arm of the five acrocentric chromosomes, (c) pericentric distribution of the two hMMT HMDPs in Chr 1, 2, 3, 4, 5, 7, 9, 10, 17 and Y, (d) the [AATGG]_n_^h^ and [CCATT]_n_^h^ hMMT HMDPs are found on the main arms of the chromosomes (on arm distribution) that are majorly in Chr 1, 2, 16, 20 and 22) (Fig. 3-E-a, Table S10.04, Fig. S3.02). Remarkably, Chr 9 is clustered a large segment of [AATGG]_n_^h^ and [CCATT]_n_^h^ hMMT HMDPs at the pericentromeric region of q-arm (Fig. 3-E-b), this region well corresponds to the heterochromatic regions consisting of Classical Human Satellite III [43, 44].

### 3.6. Comparison of HMDPs between T2T-CHM13 and GRCh38

The microsatellite density landscapes of 24 chromosomes of the reference human genome GRCh38.p14 were also built here, As the reference genome GRCh38 contained gaps and collapsed tandem repeats (Table 1-a, Table S14), the total sequence assembly of GRCh38.p14 is 2,937,639,396 bp, with unsequenced gap size of 150,630,436 bp. And only 3,480 HMDPs, 26,310 MMDPs and 178,848 LMDPs were identified in the microsatellite density landscapes of GRCh38.p14 genome (Table 1-b, Table S14); so HMDPs in GRCh38 genome are shown to be greatly different from that of T2T-CHM13, but MMDPs and LMDPs in GRCh38 genome are relatively close to those in T2T-CHM13 (Table 1-b). A detailed comparison of HMDPs in microsatellite density landscapes between full T2T-CHM13 and GRCh38 full genome, revealed that 1233 and 1290 are high and low-middle similar corresponding to those HMDPs of GRCh38 respectively (Fig. 3-F-b,c,d, Table S15, Fig. S12), above 2,000 HMDPs correspond to middle/low microsatellite density peaks (M/LMDPs) and no peaks region of GRCh38, more than half of HMDPs of T2T-CHM13 are corresponding to un-sequenced gaps region of GRCh38 (Fig. 3-F-d, Table S15); suggesting that HMDPs alleles variety possibly contribute a lot to the individual diversity of human genome in spite of that the no-sequenced gap region may influence the comparing result.

### 3.7. Genomic Regions divided by local microsatellites density features

Because the landscapes graphed by the differential calculating microsatellites density method, the STRs density distribution feature can be visualized in every 1 kb sequence local genomic region; we discovered from the landscapes that the full genome of T2T-CHM13 may be divided into 514 different microsatellites density related Genomic Regions (Table 1-d, Fig. 4-A, Fig. S4, Table S9). These Genomic Regions comprise of 4 Genomic Region class: Variable microsatellite density (V-) Region class, Even microsatellite density (E-) Region class, Peaks Cluster (PC-) Region class and Telomere repeat (T-) Region class. The V-Region class was further divided into High Variable Microsatellite Density Region (HVMD-R), Middle Variable Microsatellite Density Region (MVMD-R) and Low Variable Microsatellite Density Region (LVMD-R) subclass, the 154 Regions of this class principally correspond to gene coding regions and also some satellite distribution region (Table 1-d). The E-Region class includes relatively Even and Average Microsatellite Density Region (EAMD-R) subclass and relatively Even and extreme Low Microsatellite Density Region (EeLMD-R) subclass, 67 Regions of this class mainly match _α_ satellites and other satellites. The PC-Region class are subdivided into di-, tri-, tetra-, and penta-subclasses, Regions of this class showed that special motif type of Microsatellite Density Peaks (MDPs) usually gather in proximity to form peaks cluster. And the hexa-nucleotide motif microsatellite cluster Region class was discovered only locating in the telomeric region of T2T-CHM13, the 24 (TTAGGG)_n_-Rs were found in the telomere of q-arm of all chromosomes, as well as the reverse complementary 23 (CCCTAA)_n_-Rs were found in the telomere of p-arm of all chromosomes except the H(AATGG)_n_-R in the p-arm of Chr13 (For simplicity, the Region name abbreviation are only shown here and below, Region name nomenclature please refer to Table S9).

**Figure 4.**
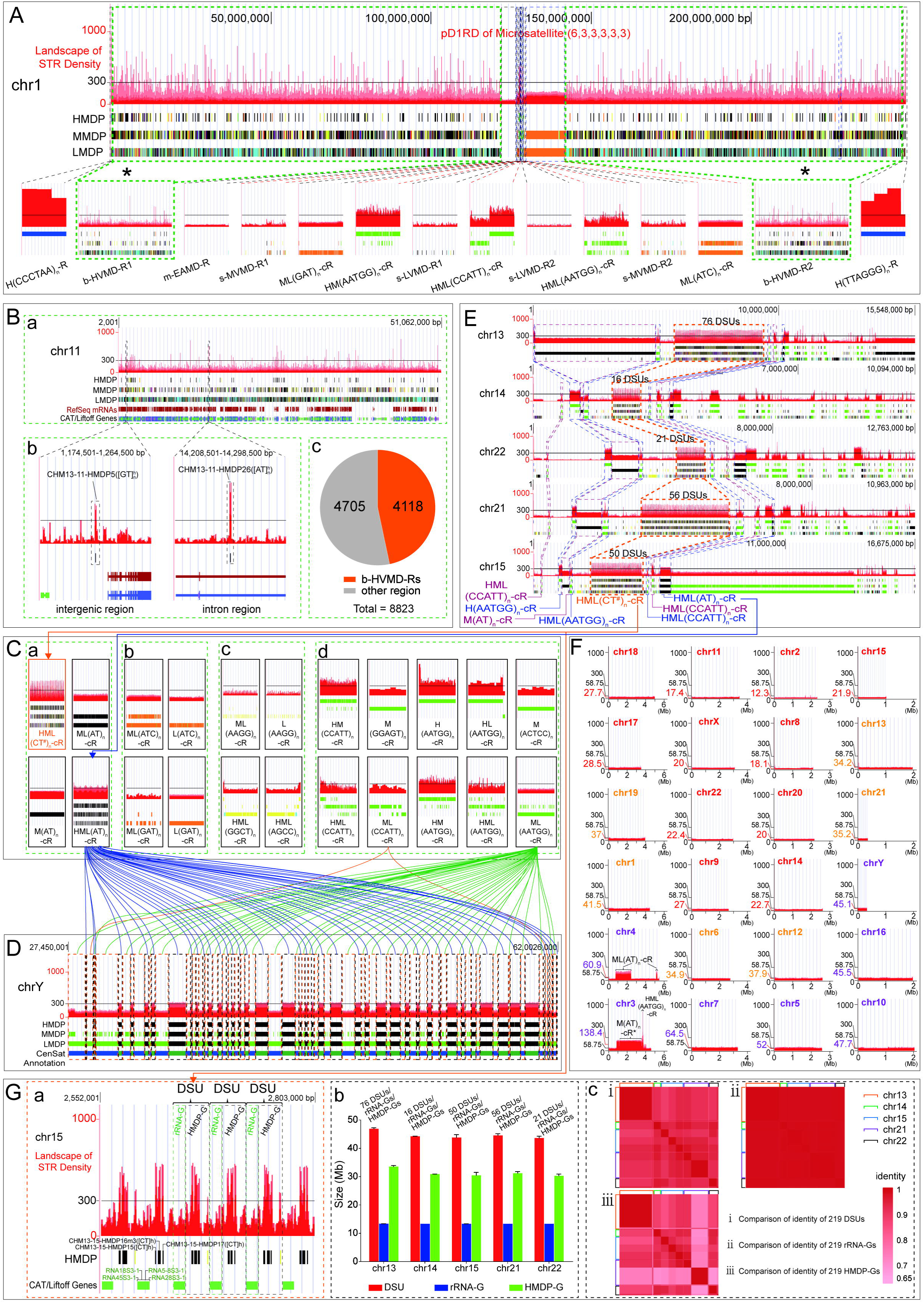
Summary of Genomic Regions characterized by microsatellite density in T2T-CHM13. **A.** Chromosome 1 of the T2T-CHM13 characterized into 14 Genomic Regions. **B.** broad High Variable Microsatellite Density Region (b-HVMD-R), (a) example of a b-HVMD-R (chromosome 11, including RefSeq mRNAs and CAT / Liftoff Genes Track). (b) HMDPs corresponding to gene intron and intergenic region; (c) the number of HMDPs in b-HVMD-Rs and other regions. **C.** Feature maps of Peaks Cluster (PC-) Regions (a: di-, b: tri-, c: tetra- and d: penta-subclass). **D.** HML(AT)_n_-cRs and ML(AATGG)_n_-cRs alternatively connecting in the Chr Y heterochromatin. **E.** Similar local microsatellites density features in the short arms of acrocentric chromosomes (HML(CT^#^)_n_-cR in center). **F.** Microsatellite density feature of centromeres. The ordinate shown average pD_1_RD value, the large font number: the average value of pD_1_RD of each centromere (colors showing different value ranges), 58.75: the average pD_1_RD of full genome. **G.** The (CT#)_n_ HMDPs groups separate rRNA genes in the five HML(CT^#^)_n_-cR, (a) Partial enlarged view of HML(CT^#^)_n_-cR in Chr15; (b) Sequence size of Duplication Segment Units (DSUs), rRNA-Groups (rRNA-Gs) and HMDP-Groups (HMDP-Gs) in HML(CT^#^)_n_-cRs; (c) Comparison of identity of DSUs, rRNA-Gs and HMDP-Gs.

### 3.8. The broad High Variable Microsatellite Density Region

The microsatellite density landscape maps display that the density of microsatellites on the main arms of most chromosomes has a huge rhythmic change, showing that HMDPs, MMDPs and LMDPs appear alternately, such regions are summarized as broad High Variable Microsatellite Density Regions (b-HVMD-Rs) with every genomic region size more than 10,000,000 bp (Fig. 4-A, Table S9). Total 42 b-HVMD-Rs were classified in the HVMD-R subclass, all chromosomes contain two b-HVMD-Rs in main central part of two arms except that the five acrocentric chromosomes include only one b-HVMD-R in each of their q-arm and ChrY include one in p-arm (Fig. 4-B-a, Table S9). Comparing all b-HVMD-Rs with CAT/Liftoff Genes, RefSeq mRNAs and CenSat Annotation tracks, demonstrated that the b-HVMD-Rs well corresponding to broad gene coding regions in every chromosome (Table 1-d, Table S9). The sum size of the 42 b-HVMD-Rs is 2,850,947,000 bp, representing 91.46% of the full sequence size of T2T-CHM13 genome; but only 4118 of the 8823 HMDPs located in the intergenic and intron region of all b-HVMD-Rs (Fig. 4-B-c, Table S10 & S11).

### 3.9. Microsatellite Density Peaks Clustering (PC-) Regions

In the 246 Genomic Regions of PC-Region class (Table 1-d, Fig. 4-C, Fig. S6-S9); we clarified 70 Regions in the di-subclass, these Regions were classified into 4 Region types as HML(CT^#^)_n_-cR, HML(AT)_n_-cR, M(AT)_n_-cR and ML(AT)_n_-cR; the 5 HML(CT^#^)_n_-cR are found in the near central part of short arm of every the five acrocentric chromosomes and correspond to the five rRNA genes coding regions; the 53 HML(AT)_n_-cR occur in the five short arm of the five acrocentric chromosomes and the heterochromatin region of ChrY; and the 3 M(AT)_n_-cR and 9 ML(AT)_n_-cR appear mainly in the five short arm of the five acrocentric chromosomes and centromeric regions of Chr 3 and 4. Secondly, 13 Regions in the tri-subclass were clarified into 4 Region types as ML(ATC)_n_-cR, L(ATC)_n_-cR, ML(GAT)_n_-cR and L(GAT)_n_-cR, which are mainly presenting at the pericentromeric regions of chr1, 2, 7, 10, 16, and 17. Then, only 5 Regions were identified in the tetra-subclass as 4 Region types as ML(AAGG)_n_-cR, L(AAGG)_n_-cR, HML(AGCC)_n_-cR and HML(GGCT)_n_-cR, occurring at Chr2, 3 and 22. At last, we sorted out 158 Regions in the penta-subclass as 10 Region types; therein 8 Region types included 156 Regions containing the reverse complementary motif of (AATGG)_n_ or (CCATT)_n_, were found at almost all chromosomes except chr6, 8, 11, 12, 16, 19, and X, and this is consistent with the observation of aforesaid no [AATGG]_n_^h^ and [CCATT]_n_^h^ motif type HMDPs distribution; the other two Region types include only two Regions also containing another two reverse complementary motif of (ACTCC)_n_ and (GGAGT)_n_, are only found at the short arm of the acrocentric chromosome 14 and 15. The comparing with CenSat Annotation tracks in UCSC genome browser reveals that di-, tri-, tetra-, and penta-subclasses mainly correspond to diverse classical human satellite sequences region (Table 1-d, Table S9).

### 3.10. HML(AT)_n_-cR and ML(AATGG)_n_-cR alternatively connecting to form the Y heterochromatin

Previously we have constructed the landscapes of the incomplete Y-DNA sequence of human genome GRCh38, which includes many unsequenced gaps especially a big gap of more than 30,000,000 bp in the heterochromatin [36]. Herein, we made the landscape of Y-DNA again but for the first complete sequence of human genome T2T-CHM13 [29], illustrating the local microsatellite relative density feature of the Y heterochromatin (Fig. 4-D), in which Genomic Region HML(AT)_n_-cR clustering of (AT)_n_ HMDPs, MMDPs and LMDPs alternatively link to Region ML(AATGG)_n_-cR clustering (AATGG)_n_ MMDPs and LMDPs; therefore, the Y heterochromatin contains 43 HML(AT)_n_-cRs and 36 ML(AATGG)_n_-cRs alternatively connected at intervals (Fig. 4-D), and only 2 ML(CCATT)_n_-cRs and 5 L(AATGG)_n_-cRs replace ML(AATGG)_n_-cRs to separate HML(AT)_n_-cRs. The above-mentioned densely distributing 2575 [AT]_n_^h^ HMDPs scatter among the 43 HML(AT)_n_-cRs and jaggedly separated by (AATGG)_n_ MMDPs and LMDPs. Comparing the landscape track with CenSat Annotation track in UCSC genome browser showed that HML(AT)_n_-cRs and ML(AATGG)_n_-cRs well correspond to alternating pattern of classical human satellite 1B (hsat1B) and classical human satellite 3 (hsat3) in the heterochromatin Yq12 region [29] (Fig. 4-D, Fig. S10).

### 3.11. Similar local microsatellites density features in short arms of the five acrocentric chromosomes

The landscapes showed that local microsatellites density features in the five short arm of the five acrocentric chromosomes is more complicate than other autosomes. A palisade arranging HML(CT^#^)_n_ cluster Region situates in central or near central part of the short arm of these chromosomes, a set of HML(CCATT)_n_-cR, H(AATGG)_n_-cR, M(AT)_n_-cR, HML(AATGG)_n_-cR array in left side, 2HML(CCATT)_n_-cR, one HML(AT)_n_-cR and several other Regions lie orderly at the right side of the central HML(CT^#^)_n_-cR; though these Regions are different in length, they arrange in similar order (Fig. 4-E, Fig. S4 & S5). Comparing with CAT/Liftoff Genes, RefSeq mRNAs and CenSat Annotation tracks, revealed that the HML(CT^#^)_n_-cR well correspond to the rRNA gene cluster Regions. Moreover, there are also other PC-Regions arraying in either side of the HML(CT^#^)_n_-cR. In a ward, the local microsatellites density features arraying in the landscapes are very similar in the five short arms of the acrocentric chromosomes.

### 3.12. Even and Low microsatellites density mainly distribute in the centromere

Although the centromeres are known as comprised of large arrays of tandem repeated alpha satellite [27] and most Genomic Regions of the T2T-CHM13 genome exhibit large variations local STRs density (ie. The pD_1_RD value) in the microsatellite density landscape maps, the centromeric regions showed that the feature of local relative microsatellite density is relatively even and low (Fig. 4-F, Fig. S10); the average pD_1_RD of microsatellite in all centromeric regions is 42 and far lower than average relative microsatellite density with value of 58.75 in the full genome, the lowest average pD_1_RD is only 12.3 in centromere of Chr2, the average pD_1_RD value of centromeres in 11 chromosomes (chr18, 11, 2, 15, 17, X, 8, 22, 20, 9, 14) are lower than 30, the average pD_1_RD value of centromeres in 6 chromosomes (chr19, 1, 6, 12, 13, 21) are between 30 to 42 (the average pD_1_RD of all centromeres), and 7 centromeres in chromosomes (chr 4, 3, 7, 5, 10, 16, Y) with the average pD_1_RD value higher than 42. Two M(AT)_n_-cRs are embedded in the landscapes of centromere in Chr 3 and 4, and cause the average pD_1_RD of centromeres rise to 138.4 and 60.9 respectively.

### 3.13. (CT^#^)_n_ HMDPs groups separate rRNA genes in the five acrocentric short arms

As aforesaid, the five short arm of the acrocentric chromosomes contain five HML(CT^#^)_n_-cRs in their central part, series HMDPs group with (CT)_n_ as main motif were found clustering palisadingly and followed by several MMDPs and LMDPs, forming a tandem Duplication Segment Unit (DSU), these DSUs connects tandemly composing these HML(CT^#^)_n_-cRs. The size of these DSUs are between 43000-47000 bp considered as rDNA array [24], and there are 76 DSUs found in the short arm of Chr13, 16 DSUs in Chr14, 50 DSUs in Chr15, 56 DSUs in Chr21, 21 DSUs in Chr22, and total 219 DSUs. Further comparison show that every HML(CT^#^)_n_-cR comprise of the rRNA genes Groups region and the [CT^#^]_n_ HMDPs Groups region, remarkably, the [CT^#^]_n_ HMDPs Groups region well one-by-one separate every the rRNA genes Group in every DSU of the five HML(CT)_n_-cR Regions (Fig. 4-G-a, Fig. S5). Alignment results showed that the identities among the 219 DSUs, 219 rRNA genes Groups and 219 HMDPs Groups are higher than 0.82, 0.99 and 0.68 respectively (Fig. 4-G-b & c, Fig. S5), suggesting the [CT^#^]_n_ HMDPs Groups region may functionally separate the rRNA genes region but be more variable than rRNA genes region.

## 4. Discussion

This work built the first comprehensive genome wide microsatellite density landscape maps of the first complete human reference genome T2T-CHM13 [24, 29]. These landscape maps exhibited that microsatellites aggregate in many genomic positions to form a large number of microsatellite density peaks, and these peaks array along the every human chromosomes arranged like notes of a beautiful piece of music; therefore, the 24 microsatellite density landscapes together look like to form a symphony of human life (Fig. 2). Though transcriptional and epigenetic state of human repeat elements were comprehensive analyzed [5], we mainly focus on the short tandem repeats of human genome, and our works revealed that the microsatellite density landscapes are compatible with human chromosome structure at a high extent, suggesting these landscapes possibly contribute to the further exploration the relationship between STRs aggregation and human genome structure, variation, evolution and so on.

The high microsatellites density peak (HMDP) aggregated predominantly by a single motif, is the most statistically significant short tandem repeats aggregation phenomenon. We identified 8823 HMDPs in T2T-CHM13, much more than the number of 3480 HMDPs in GRCh38; the HMDP number was substantially increased in T2T-CHM13, mainly corresponding to unassembled regions in GRCh38, especially the heterochromatin of Chr Y and the five short arms of the acrocentric Chromosomes. Almost all the HMDPs are single motif dominated, particularly, 86.1% of the 8823 HMDPs are high percentage dominated by a single motif. And there should be total 964 Main Motif Type of HMDPs ranging from mono- to hexa-main motif types, actually, we only identified 87 single motif dominated Main Motif Types (MMTs); these statistical significances are strongly implied the importance of aggregation of short tandem repeats on gene expression, regulation, epigenetics, genetic architecture and evolution of human genome. What is more, the (AT)_n_ single motif dominated Main Motif Types (MMTs) account for more than half of the total number of HMDPs, and the reverse complementary motif (AATGG)_n_ and (CCATT)_n_ dominated two Main Motif Types (MMTs) account for approximately half of the total bins of HMDPs; suggesting that the three MMT HMDPs are worthy to be further explored for their roles in gene function and chromosome structure of human genome.

In addition, we classified 514 characterized by microsatellite density in the complete reference genome, the Variable microsatellite density (V-) region class almost overlap the gene coding region, the Even microsatellite density (E-) region class mainly cover centromere, pericentromere and p-arm of acrocentric chromosomes, especially, the m-EAMD-R and m-EeLMD-R well correspond to centromeres (Table S9.01); the 246 PC-regions generally locate in pericentromere, p-arm of acrocentric chromosomes and heterochromatin of Chr Y; and the reverse complementary repeat motif (CCCTAA)_n_ and (TTAGGG)_n_ telomere repeat regions almost dominate in p-arms and q-arms respectively. Meanwhile, similar genomic regions structural characters were observed in the five acrocentric short arms by the view of microsatellite density features, and alternative microsatellite density regions structure observed in heterochromatin Chr Y [29]. all these indicate that local regional microsatellite density variation may be related to human genome structure. Furthermore, most (microsatellite density) peaks cluster (PC-) regions correspond to different Classical human satellites (hsat); but most the even microsatellite density regions especially in centromere, where microsatellite density are limited to very low and even, correspond to different α satellite higher-order repeats (αSat hor) (Table 1-d); suggesting that although microsatellites (STRs) are highly related to satellites (long tandem repeats), the mechanism of their occurrence are likely completely different.

The 219 near identical Duplication Segment Units (DSUs) of approximately 45-kbp that contain segments of encoding 45S rRNAs, are embedded in the short arms of the 5 acrocentric chromosomes [5]; we identified that the HMDPs also well separate every 45S rRNA gene of about 13300 bp. All the HMDPs were classified here in the intergenic and intron regions, implying that the HMDPs possibly own the basic role of separating genes and exons in the genome and may be very worthwhile to further explore. The first complete reference T2T-CHM13 provided us the opportunity to study short tandem repeat sequences in human genome comprehensively, our results of microsatellite density landscape maps revealed that short tandem repeats are tend to aggregate characteristically throughout the full genome of T2T-CHM13, it may be very helpful to deepen the exploring of the mysterious roles of tandem repeats to human genome’s structure, evolution, regulation, variation and also human diseases.

## 5. Conclusions

These landscape maps exhibited that microsatellites aggregate in many genomic positions to form a large number of microsatellite density peaks with composing of mainly single motif type in the complete reference genome, indicating that the local microsatellites density varies enormously along the every chromosome of T2T-CHM13.

## Declarations

### Abbreviations

STRs: Short Tandem Repeats
HMDPs: High Microsatellites Density Peaks
MMDPs: Middle Microsatellites Density Peaks
LMDPs: Low Microsatellites Density Peaks
MDPs: Microsatellite Density Peaks
DCM 2.0: Differential Calculator of Microsatellites version 2.0
IMEx 2.1: Imperfect Microsatellite Extractor version 2.1
pD_1_RD: position-related D_1_-Relative Density
MDPS.v1.0: Microsatellite Density Peaks Sorter version 1.0
MMT: Main Motif Type
sub-MMT: sub-Main Motif Type
GHC.v1.0: Genome HMDPs Comparator version 1.0
RD: Relative Density
hMMT: high-percentage Main Motif Type
mMMT: middle-percentage Main Motif Type
lMMT: low-percentage Main Motif Type
AD: Average Distance
V-region class: Variable microsatellite density region class
E-region class: Even microsatellite density region class
PC-region class: Peaks Cluster region class
T-region class: Telomere repeat region class
HVMD-R: High Variable Microsatellite Density Region
MVMD-R: Middle Variable Microsatellite Density Region
LVMD-R: Low Variable Microsatellite Density Region
EAMD-R: Even and Average Microsatellite Density Region
EeLMD-R: Even and extreme Low Microsatellite Density Region
DSUs: Duplication Segment Units
b-HVMD-Rs: broad High Variable Microsatellite Density Regions
hsat: human satellites
αSat hor: α satellite higher-order repeats

### Ethics approval and consent to participate

Not applicable.

### Consent for publication

Not applicable.

### Availability of data and materials

The Web version of microsatellite density landscape maps of 24 chromosomes of T2T-CHM13v2.0 can be accessed in (http://genome.ucsc.edu/s/zhongyangtan/CHM13v2.0).

The Web version of microsatellite density landscape maps of 24 chromosomes of GRCh38.p14 can be accessed in (http://genome.ucsc.edu/s/zhongyangtan/GRCh38.p14).

All of the Supplementary Figures and Tables can be accessed in https://drive.google.com/file/d/1DiXYuUnSINpdkfCbZJAyMkPBKIFfwFEg/view?usp=sharing

### Authors’ contributions

Z.T. designed and directed this study. Z.T., K.O., Y.X., D.L., T.C., Sc.P., H.H., W.Z. and Y.L. prepared the manuscript. T.C., D.L., Sc.P., H.H., Y.F., W.Z. and Z.T. developed the statistical method and programs. Y.X., D.L., T.C., Sc.P., H.H., W.Z., Y.L., Z.P., H.Z., Li.Z., S.P., R.S., X.H., S.Z., W.J., X.Z., X.W., La.Z., J.Z., Q.O., Y.T., X.J., Y.Z., S.T., J.S. and Z.T. performed the data analysis, K.O. and Z.T. edited this manuscript. All authors read and approved the final version of the manuscript.

## Acknowledgments

We thank Yongliang Wu (The Hong Kong Polytechnic University), Jingchen Xie, Junyu Zeng and Wenjing Zhou (Hunan University), Yasha He (South Central University), Puxi Tan (Xi’an Jiaotong University) for helping in data analysis.

